# Using paired serology and surveillance data to quantify dengue transmission and control during a large outbreak in Fiji

**DOI:** 10.1101/246116

**Authors:** Adam J Kucharski, Mike Kama, Conall H Watson, Maite Aubry, Sebastian Funk, Alasdair D Henderson, Oliver J Brady, Jessica Vanhomwegen, Jean-Claude Manuguerra, Colleen L Lau, W John Edmunds, John Aaskov, Eric J Nilles, Van-Mai Cao-Lormeau, Stephane Hue, Martin Hibberd

## Abstract

Dengue is a major health burden, but it can be challenging to examine transmission dynamics and evaluate control measures because outbreaks depend on multiple factors, including human population structure, prior immunity and climate. We combined population-representative paired sera collected before and after the major 2013/14 dengue-3 outbreak in Fiji with surveillance data to determine how such factors influence dengue virus transmission and control in island settings. Our results suggested the 10-19 year-old age group had the highest risk of acquiring infection, but we did not find strong evidence that other demographic or environmental risk factors were linked to seroconversion. Mathematical modelling showed that temperature-driven variation in transmission and herd immunity could not fully explain observed dynamics. However, there was evidence of an additional reduction in transmission coinciding with a vector clean-up campaign, which may have contributed to the decline in cases and prevented transmission continuing into the following season.

## Introduction

In recent years, the incidence of dengue has risen rapidly. In the Asia-Pacific region, which bears 75% of the global dengue disease burden, there are more than 1.8 billion people at risk of infection with dengue viruses (DENV) [1]. Increased air travel and urbanisation could have contributed to the geographic spread of infection [2, 3], with transmission by mosquitoes of the *Aedes* genus, including *Aedes aegypti* and *Aedes albopictus* [4]. DENV has four serotypes circulating, with infection conferring lifelong protection against the infecting serotype and short-lived protection against the others [5, 6]. Although four serotypes of DENV may co-circulate in South East Asia, only one serotype circulates in most of the South Pacific islands at any point in time [7, 8].

Between November 2013 and July 2014, a major outbreak caused by DENV-3 occurred in Fiji, with more than 25,000 suspected cases reported (Figure 1A). Prior to the 2013/14 outbreak, there were eleven outbreaks of dengue recorded in Fiji, involving serotypes 1, 2 and 4 (Table 1). Most cases in 2013/14 occurred on Viti Levu, the largest and most populous island. This is administratively divided into the Central Division, which includes the port-capital Suva, and Western Division, which contains the urban centres of Lautoka and Nadi, where Fiji’s major international airport is located. Dengue transmission in Central and Western Divisions is likely to be driven mostly by the *Aedes aegypti* vector, with *Aedes albopictus* most abundant in the Northern Division. *Aedes polynesiensis* and *Aedes pseudoscutellaris* are also present in all divisions [9, 10]. In response to the 2013/14 outbreak, considerable resources were dedicated to implementing control measures, including a nationwide vector clean-up campaign between 8th and 22nd March 2014 [11]. As well as media coverage and distribution of flyers to raise awareness about dengue prevention and protection, a major operation was put in place to remove rubbish that could act as egg laying habitats for mosquitoes. In total, forty-five tonnes of tyres and twenty-five tonnes of other containers were removed during this period.

**Figure 1:**
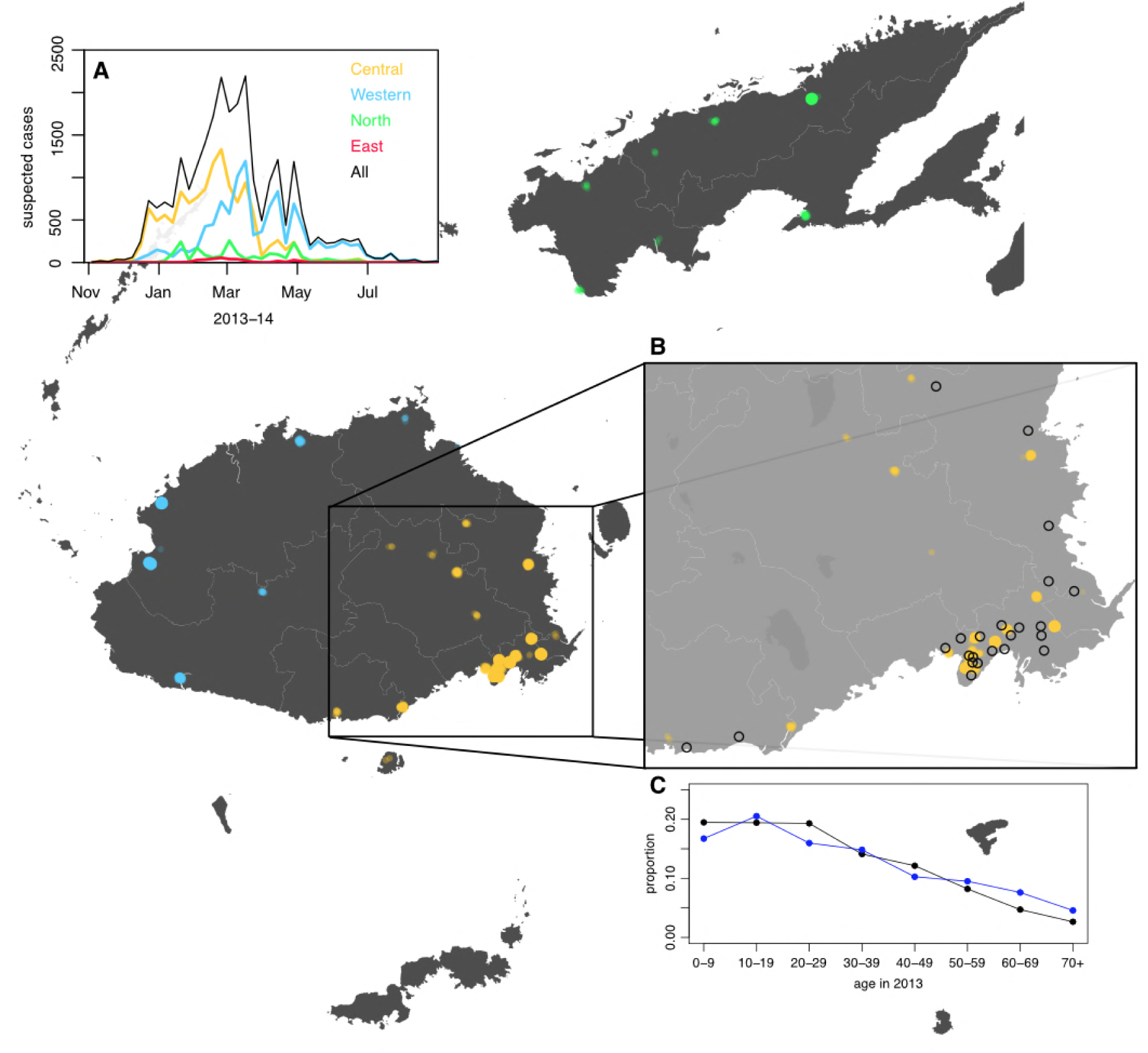
Geographical distribution of suspected dengue cases in Northern (green), Western (blue) and Central (yellow) divisions between 27th October 2013 and 1st July 2014. Points on the maps show locations of cases arranged by health centre they reported to; these are plotted with jitter and transparency to show concentrations of cases. (A) Weekly reported case totals for Northern, Western, Central and Eastern divisions. (B) Serosurvey study locations. Black circles show the 23 study clusters included in the analysis. (C) Age distribution of Central Division in the 2007 census (blue line) and ages of serosurvey participants in 2013 (black line).

**Table 1:** Reported dengue outbreaks in Fiji between 1930–2014. Two studies [22, 12] also included a post-outbreak serosurvey in Central Division.

| Year | Serotype | Reported cases | Seroprevalence | Source |
| --- | --- | --- | --- | --- |
| 1930 | ? | Thousands |  | [9] |
| 1944-5 | 1 | Thousands |  | [53] |
| 1971-3 | 2 | 3,413 | 26% (Suva) | [22] |
| 1974-5 | 1 | 16,203 |  | [53] |
| 1980 | 4 | 127 |  | [12] |
| 1981 | 1 | 18 |  | [54] |
| 1982 | 2 | 676 |  | [54] |
| 1984-6 | ? | 490 |  | [12] |
| 1988 | ? | 22 |  | [12] |
| 1989-90 | 1 | 3,686 | 54% (Suva) | [12, 23] |
| 1997-8 | 2 | 24,780 |  | [55] |
| 2001-3 | 1 | ? |  | [56] |
| 2008 | 4 | 1,306 |  | [57, 58] |
| 2013-14 | 3 | 25,496 |  | Fiji MOH |

Large dengue outbreaks can place a substantial public health burden on island populations [12, 13]. However, understanding the dynamics of infection and evaluating the impact of vector control measures remains challenging. There is a limited evidence base for control measures even in controlled trials [14, 15], and post-outbreak evaluation is hindered by the fact that the size and duration of major outbreaks can be influenced by several factors, including population immunity, human movement, seasonal variation in transmission, and proportion of people living in urban, peri-urban and rural communities. In Fiji, dengue outbreaks typically occur during the wetter, warmer season between December and July, when vectors are most abundant [16]. Although surveillance data can provide broad insights into arbovirus transmission patterns [17, 18, 19], and cross-sectional sero-surveys can be used to measure contemporary levels of immunity [20, 21, 22, 23], characterising infection dynamics in detail requires cohort-based seroepidemiological studies [24, 25], which can be difficult to implement in island settings where outbreaks are infrequent and difficult to predict.

Immediately before the 2013/14 dengue outbreak in Fiji, a population-representative serological survey had been conducted to study leptospirosis and typhoid [26]. To investigate patterns of dengue infection in 2013/14, we followed up participants from this survey in Central Division, to obtain a set of paired pre- and post-outbreak serological samples. We tested the paired samples for anti-DENV IgG antibodies using ELISA and a recombinant antigen-based microsphere immunoassay (MIA), and combined these data with dengue surveillance data to compare possible explanations for the outbreak dynamics. We measured age-specific and spatial patterns of infection and reported disease, and tested whether there were demographic and environmental risk factors associated with infection. Having characterised factors shaping individual-level infection risk, we used a Bayesian approach to fit a transmission dynamic model to both the serological survey and surveillance data in order to estimate the contribution of climate and control measures to the decline in transmission observed in 2014.

## Materials and methods

### Surveillance data

In December 2013, the dengue outbreak in Fiji was determined to be due to DENV-3 by RT-PCR performed on serum samples sent to the World Health Organization Collaborating Centre for Arbovirus Reference and Research at the Queensland University of Technology (QUT, Brisbane). Hereafter, samples that were ELISA reactive for NS1 antigen or IgM were presumed to be to DENV-3 infection with a sub-sample of subsequent positive samples sent for confirmatory serotyping at QUT, the Institut Louis Malardé (ILM) and the US Centers for Disease Control and Prevention. Of the 10,442 laboratory tested cases that were notified to the Fiji National Centre for Communicable Disease Control between 27th October 2013 and 4th March 2014, 4,115 (39.4%) were reactive for DENV NS1 and/or anti-DENV IgM (Figure S1). After this time period, dengue surveillance was transitioned from laboratory to clinical-based reporting (i.e. dengue-like illness, DLI) due to the size of the outbreak (Figure S1).

Between 27th October 2013 and 31st August 2014, 25,494 suspected cases of dengue (i.e. laboratory tested or confirmed or DLI) were notified to the Fijian Ministry of Health. Of these, 12,413 (48.7%) cases were in Central Division, predominantly in the greater Suva area (Figure 1). 10,679 cases were reported in the Western Division, 2,048 cases were reported in the Northern division, largely in or near Labasa, the largest town of Vanua Levu island, and 354 cases were reported in the Eastern Division. For the lab confirmed cases, date of testing was used to compile weekly case incidence time series; for the DLI data, date of presentation to a health centre was used, as these dates were most complete. Filter paper-based surveillance conducted by ILM between December 2013 and October 2014 found 24 samples positive for DENV-3 by RT-PCR, as well as three samples positive for DENV-2 and one for DENV-1. During 2014/15, there was a flare up of DENV-2 in Fiji. However, relatively few cases occurred on Viti Levu: of the 543 confirmed cases nationally between 1st January 2015 and 29th April 2015, 437 cases (80%) were from the Northern Division [27].

### Serological survey

We conducted a serological survey using pre- and post-outbreak sera from 23 communities in Central Division. Pre-outbreak sera were collected as part of population representative community-based surveys of leptospirosis and typhoid conducted in Central Division between September and November 2013 [26, 28]. Population-proportionate sampling was used to select local nursing zones (the smallest administrative unit). From each of these zones, one community was randomly selected, followed by 25 households from each community and one individual from each of the households. Coincidentally, the sample collection in Central Division finished the same week as the first dengue cases were reported (Figure S2). Post-outbreak sera were collected during a follow-up study carried out in October and November 2015. Field teams visited participants in Central Division who had previously participated in the 2013 serological study and had consented to being contacted again for health research.

Participants who gave informed consent for the 2015 study completed a questionnaire and provided a 5ml blood sample. The study was powered to measure the rise in prevalence of anti-DENV antibodies between 2013–15. Historical dengue outbreaks in Fiji (Table 1) suggested we would expect to see seroconversion in at least 20% of the study population. Allowing for 5% seroreversion, and 0.05 probability of type-1 error, McNemar’s test suggested 250 paired samples could detect a 15% increase in seroprevalence with 95% power, and a 20% increase with ≈100% power. We also collected data on potential risk factors and healthcare-seeking behaviour during this period. The questionnaire asked for details of fever and related visits to a doctor in the preceding two years, and the same for household members in the preceding two years. The questionnaire also recorded details of household environment, including potential mosquito breeding grounds (File S1).

### Ethical considerations

The 2013 typhoid and leptospirosis studies and the 2015 follow up study were approved by the Fiji National Research Ethics Review Committee (ref 2013-03 and 2015.111.C.D) and the London School of Hygiene & Tropical Medicine Observational Research Ethics Committee (ref 6344 and 10207). Participants in the 2015 follow up study were people who had previously given informed consent to have their blood tested as part of a public health serum bank established in the 2013 typhoid and leptospirosis serosurvey, and agreed to be contacted again by public health researchers. The study was explained in English or the local iTaukei language by bilingual field officers, at the potential participants’ preference. Adults gave written informed consent, or thumbprinted informed consent witnessed by a literate adult independent from the study. For children age 12–17 years, written consent was obtained from both the parent and the child. For children aged under 12 years, written consent was obtained from the parent only, though information was provided to both.

### Serological testing of paired sera

Paired pre- and post-outbreak serum samples were tested using an indirect IgG ELISA kit (PanBio Cat No 01PE30), according to manufacturer guidelines. This assay employs recombinant DENV envelope proteins of all four serotypes [29]. Samples with ELISA value of ≤9 PanBio units were defined as seronegative, ≥11 PanBio units seropositive, and values between 9 and 11 as equivocal. Seroconversion was defined as a change from seronegative to seropositive status. Samples were also tested against each of the four specific DENV serotypes using a recombinant antigen-based microsphere immunoassay (MIA) [20, 30]. Defining a reactive MIA result against at least one serotype to be DENV seropositive, 80% of participants had the same DENV serostatus (i.e. both seropositive or both seronegative) by ELISA and MIA (Cohen’s *κ*=0.52).

### Serological modelling

To estimate the probability that a given increase in ELISA value was the result of a genuine rise rather than measurement error, we fitted a two distribution mixture model to the distribution of changes in value between 2013 and 2015. We used a normal distribution with mean equal to zero to reflect measurement error, and a gamma distribution to capture a rise that could not be explained by the symmetric error function. The observed changes in ELISA value we fitted to ranged from −6 to 20; we omitted the two outliers that had a change in value of −9 between 2013 and 2015, as these could not be explained with a normal measurement error function. We used a generalized additive model with binomial link function to examine the relationship between value in 2013 and rise between 2013 and 2015, with data points weighted by probability of infection. Risk factor analysis was performed using a univariable logistic regression model. Both were implemented using the mgcv package in R version 3.3.1 [31, 32].

### Transmission model

We modelled DENV transmission dynamics using an age-structured deterministic compartmental model for human and vector populations, with transitions between compartments following a susceptible-exposed-infective-removed (SEIR) structure [33, 34, 35]. As human population size was known, but the vector population was not, the human compartments were specified in terms of numbers and vectors in terms of proportions. Upon exposure to infection, initially susceptible humans (*S_h_*) transitioned to a latent class (*E_h_*), then an infectious class (*I_h_*) and finally a recovered and immune class (*R_h_*). The mosquito population was divided into three classes: susceptible (*S_v_*), latent (*E_v_*), and infectious (*I_v_*). Mosquitoes were assumed to be infectious until they died. We had two human age groups in the model: aged under 20 (denoted with subscript *c*), and aged 20 and over (denoted with subscript *a*). We included births and deaths for the vector population, but omitted human births and deaths because the mean human lifespan is much longer than the duration of the outbreak. The model was as follows:

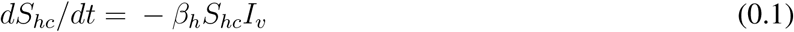

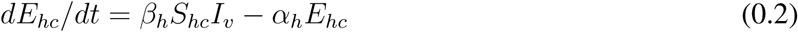

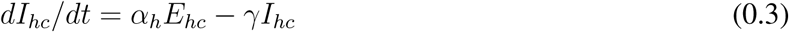

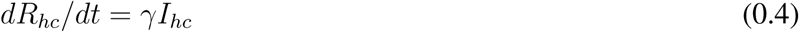

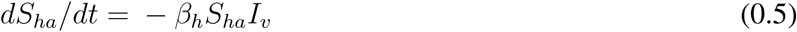

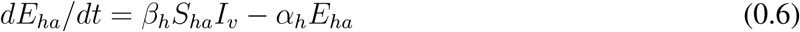

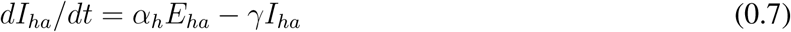

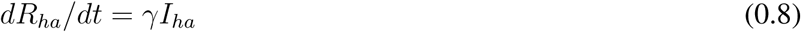

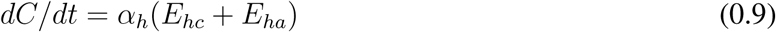

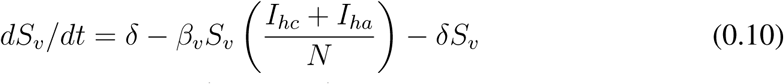

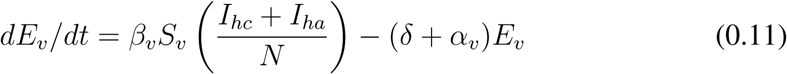

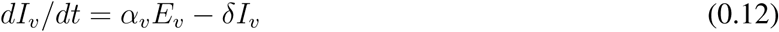

The compartment *C* recorded the cumulative total number of human infections, which was used for model fitting. Parameter definitions and prior distributions are given in Table S2. We used informative priors for the extrinsic latent period, 1/*α_v_*, mosquito lifespan, 1/*μ*, intrinsic latent period, 1/*α_H_*, and human infectious period, 1/*γ*. Based on most recent Fiji census in 2007, we set the population size *N* to be 342,000 in Central Division [36], and split this population between the two age groups based on the populations of each reported in the census (*N_c_*=133,020 and *N_a_*=208,980). We estimated two initial conditions for each human age group: the initial number of infective individuals, 
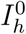
, and the initial number immune, 
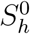
 We assumed that there were the same number of individuals initially exposed as there are individuals infectious 
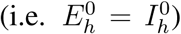
. For the vector population, we only estimated the initial proportion infectious. We assumed that 
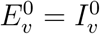
 and the remaining proportion of mosquitoes were susceptible.

We assumed the vector-to-human and human-to-vector transmission rates (*β_h_* and *β_v_*) could potentially vary over time in two ways in the model. First, transmission could fluctuate due to seasonal changes in temperature and rainfall [37]. During 2013/14 in Central Division, average monthly rainfall ranged from around 100 to 400mm, and temperature between 22 and 26°C [38]. Temperature reached its maximum in February, and minimum in August/September (Figure S3A). There is evidence that dengue transmission increases monotonically with temperature up to 30°C [39]. We therefore used a sinusoidal function to model the seasonal impact on transmission at time *t*, which was translated by a parameter *k_season_* so that it reached its maximum on 15th February:

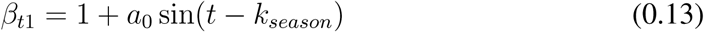

We imposed a prior distribution on the amplitude of seasonal forcing, *a*_0_: using the estimated mechanistic relationship between temperature and relative basic reproduction number *R*_0_ for arboviruses transmitted by *Aedes aegypti* [39], we calculated the relative reduction that would be expected in Fiji based on the maximum and minimum temperates, and used this value as the mean of the prior (Figure S3B–C). The relative transmission at the minimum temperature was 0.256 of the value at the maximum, which implies a mean value of *a*_0_ = (1 − 256)/(1 + 0.256) = 0.592. To avoid infection declining to implausibly small levels then rising again in the following season, we included potential for extinction in the model. If the number of individuals in any of the *E* or *I* human compartments dropped below one, the model set the value to zero. Hence if there were no exposed or infectious individuals in either of the age groups, the epidemic would end.

Second, we used a flexible sigmoid function to capture potential additional reduction in transmission over time as a result of the national clean-up campaign between 8th and 22nd March 2014:

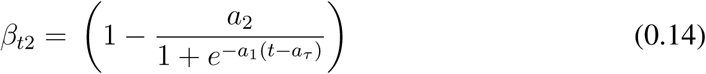

We constrained this function so that the midpoint, *a_τ_*, was between the start date of the campaign, 8th March 2014, and 5th April 2014, four weeks later (Figure S3D). We assumed that vector-to-human transmission, *β_h_*, was the product of these two time-varying functions, scaled by a baseline transmission rate, *β*_0_, i.e. *β_h_* = *β*_0_*β*_*t*1_*β*_*t*2_. The human-to-vector transmission rate was equal to this rate multiplied by a scaling factor, *a*_*v*_, to reflect potential asymmetry in transmission between humans and vectors [40]: *β*_*v*_ = *β*_*h*_*a*_*v*_. The next generation matrix for humans and vectors was defined as follows [34]:

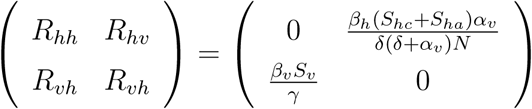

and the effective reproduction number, *R*, was equal to the dominant eigenvalue of this matrix.

### Model fitting

The model was jointly fitted to laboratory-confirmed case data and serological data using Markov chain Monte Carlo (MCMC) via a Metropolis-Hastings algorithm. For the case data, we considered time units of one week. To construct a likelihood for the observed cases, we defined case count for week *t* as *c*_*t*_ = *C*_*t*_ − *C*_*t*−1_. We assumed that observed cases followed a negative binomial distribution with mean *rc_t_* and dispersion parameter *ϕ*, to account for potential temporal variability in reporting [41]. Because reporting switched from lab confirmation to DLI during the outbreak, we fitted two sets of time series data, each with its own *r* and *ϕ* parameter. The first dataset was lab confirmed cases. We defined the first observation as 4th November 2013, the week of the first confirmed case in Central Division, and the last observation as 4th March 2014, after which reporting switched to DLI. Because there was a period of changeover, we omitted the following two weeks of data, then fitted to total suspected cases (i.e. DLI and lab tested) from 18th March until 28th July, when there had been three subsequent weeks with no reported cases. We varied the number of weeks omitted from 2–4 as a sensitivity analysis.

We also fitted the model to the proportion of each age group immune (as measured by seroprevalence) at the start and end of the outbreak. Let *X_ij_* be a binomially distributed random variable with size equal to the sample size in group *i* and probability equal to the model predicted immunity in year *j*, and *z_ij_* be the observed seroprevalence in group *i* in year *j*. The overall log-likelihood for parameter set *θ* given case data 
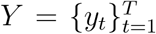
 and serological data *Z* = {*z*_*ij*_}_*i*∈{1,2},*j*∈{2013,2015}_ was therefore:

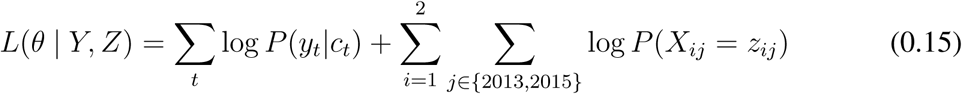

All observations were given equal weight in the model fitting. The joint posterior distribution of the parameter set *θ* was obtained from 100,000 MCMC iterations, each with a burn-in period of 10,000 iterations. We used adaptive MCMC to improve efficiency of mixing: we iteratively adjusted the magnitude of the covariance matrix used to resample *θ* to obtain a target acceptance rate of 0.234 [42]. Posterior estimates are shown in Figure S4 and correlation plots for the transmission rate parameters are shown in Figure S5. The statistical and mathematical models were implemented in R version 3.3.1 [32] using the deSolve package [43] and parallelised using the doMC library [44]. Code and data are available at: https://github.com/adamkucharski/fiji-denv3-2014

## Results

The pre- and post-outbreak serological survey included 263 participants from the Central Division, with age distribution of these participants consistent with the population distribution (Figure 1C). We found that 58.6% of participants (154/263) were ELISA seropositive to at least one DENV serotype in late 2013. Two years later, in October/November 2015, this had risen to 74.5% (196/263). Additional serotype-specific MIA tests confirmed that the largest rise in seroprevalence in Central Division was against DENV-3, from 32.7% to 52.9% (Table 2), consistent with the majority of RT-PCR-confirmed samples during the outbreak being of this serotype.

**Table 2:** Number of participants who were seropositive to DENV in 2013 and 2015 as measured by ELISA and MIA.

| Test | N | 2013 | 2013 (%) | 2015 | 2015 (%) | Difference |
| --- | --- | --- | --- | --- | --- | --- |
| ELISA | 263 | 154 | 58.6% (52.3-64.6%) | 196 | 74.5% (68.8-79.7%) | 16% (11.8-21%) |
| MIA DENV-1 | 263 | 177 | 67.3% (61.3-72.9%) | 198 | 75.3% (69.6-80.4%) | 7.98% (5.01-11.9%) |
| MIA DENV-2 | 263 | 33 | 12.5% (8.8-17.2%) | 41 | 15.6% (11.4-20.5%) | 3.04% (1.32-5.91%) |
| MIA DENV-3 | 263 | 87 | 33.1% (27.4-39.1%) | 140 | 53.2% (47-59.4%) | 20.2% (15.5-25.5%) |
| MIA DENV-4 | 263 | 79 | 30.0% (24.6-36%) | 99 | 37.6% (31.8-43.8%) | 7.6% (4.71-11.5%) |

To characterise patterns of infection between 2013 and 2015, we first considered individual-level demographic, behavioural and environmental factors. Using a univariable logistic regression model, we compared seroconversion determined by ELISA with questionnaire responses about household environment and health-seeking behaviour (Table 3). The factors most strongly associated with seroconversion between 2013–15 among initially seronegative participants were: living in an urban or peri-urban environment (odds ratio 2.18 [95% CI: 0.953-5.11], p=0.068); reporting fever in preceding two years (odds 2.94 [1.08-8.38], p=0.037); and visiting a doctor with fever in the preceding two years (odds 3.15 [1.06-10.10], p=0.043). Of the participants who seroconverted, 10/38 (26.3% [13.4-43.1%]) reported visiting a doctor with fever in the preceding two years, 2/38 (5.26% [0.644-17.7%]) reported fever but did not visit a doctor, and 26/38 (68.4% [51.3-82.5%]) did not report fever (Table S1).

**Table 3:** Risk factors from a univariable logistic regression model. Sample population was all individuals who were seronegative in 2013 (n=97), and outcome was defined as seroconversion as measured by ELISA. Number indicates total individuals with a given characteristic.

| Variable | Number | Odds ratio | p-value |
| --- | --- | --- | --- |
| **Demographic characteristics** |  |  |  |
| Age under 18 | 59 | 0.48 (0.2-1.09) | 0.08 |
| Female | 48 | 0.81 (0.36-1.84) | 0.62 |
| iTaukei ethnicity | 85 | 1.33 (0.39-5.32) | 0.66 |
| **Environmental factors present** |  |  |  |
| Mosquitoes | 90 | 4.19 (0.68-80.85) | 0.19 |
| Used car tires | 61 | 1.80 (0.77-4.42) | 0.18 |
| Open water container(s) | 61 | 1.49 (0.64-3.58) | 0.37 |
| Air conditioning | 23 | 0.46 (0.15-1.26) | 0.15 |
| Blocked drains | 53 | 1.04 (0.46-2.38) | 0.92 |
| **Location** |  |  |  |
| Urban or peri-urban | 50 | 2.18 (0.95-5.11) | 0.07 |
| **Health seeking behaviour** |  |  |  |
| Fever in preceding 2 years | 20 | 2.94 (1.08-8.38) | 0.04 |
| Visited doctor with fever in preceding 2 years | 16 | 3.15 (1.06-10.13) | 0.04 |
| Household member visited doctor with fever in preceding 2 years | 9 | 2.08 (0.52-8.94) | 0.30 |

As well as estimating infection by measuring seroconversion based on threshold values, we also considered the distribution of ELISA values. There was a noticeable right shift in this distribution between 2013 and 2015, with ELISA values increasing across a range of values (Figure 2A). As some of the individual-level changes in value between the two tests were likely to be due to measurement error [45], we fitted a mixture model to the distribution of changes in ELISA value (Figure 2B). We used a normal distribution with mean zero to capture measurement error, and a gamma distribution to fit rise that could not be explained by this error function. The fitted model suggested that a rise in value of at least 3 was more likely to be a genuine increase rather than measurement error, as shown by the dashed line in Figure 2B.

**Figure 2:**
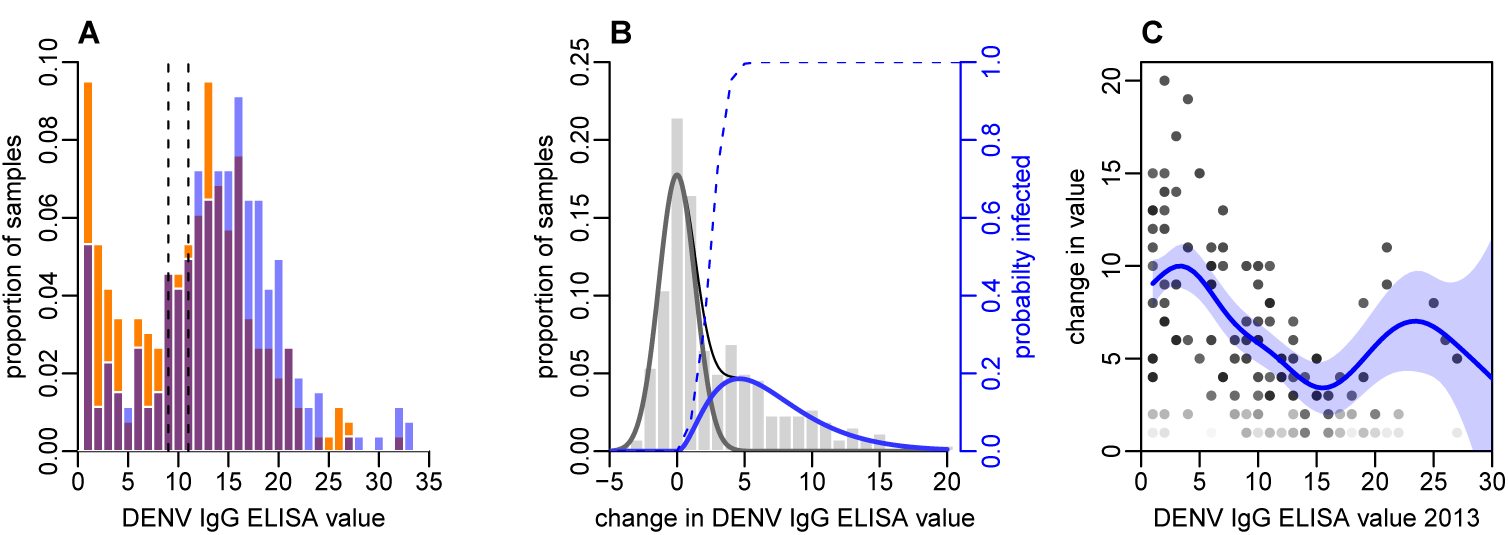
Distribution of ELISA values for anti-DENV IgG over time. (A) Distribution of values in 2013 and 2015. Orange bars show observed proportion of samples with each value in 2013; blue bars show proportions in 2015. Dashed lines show threshold for seronegativity and seropositivity. (B) Change in ELISA values between 2013 and 2015. Bars show distribution of values. Grey line shows estimated uncertainty in assay measurements; blue line shows estimated increase in value following the 2013-14 epidemic; thin black line shows overall fitted distribution (model *R*^2^=0.93). Dashed line shows probability of infection for a given rise in value. (C) Relationship between value in 2013 and rise between 2013 and 2015, adjusting for probability of infection as shown in Figure 2B. Points show 1000 bootstrap samples of the data with replacement, with opacity of each point proportional to probability of infection. Blue line shows prediction from generalized additive model, with data points weighted by probability of infection; shaded region shows 95% CI (model *R*^2^=0.31).

To explore the relationship between the initial ELISA value and rise post-outbreak, given that an individual had been infected, we fitted a generalized additive model to the data and weighted each observation by the probability that a specific participant had been infected based on the dashed line in Figure 2B. By adjusting to focus on likely infections, we found a negative relationship between initial value and subsequent rise, with ELISA values near zero rising by around 10 units, but higher values exhibiting a smaller rise (Figure 2C). Using this approach, we also found strong evidence that self-reported symptoms were associated with larger rise in ELISA value, given likely infection. Using a logistic model with self-reported symptoms as outcome and change in value as dependent variable, adjusting for initial value and again weighting by probability of infection, we found that individuals who reported a fever in the preceding two years had a predicted rise in ELISA value that was 2.2 (95% 0.77-3.6) units higher than those who did not (p=0.003). Further, individuals who reported visiting a doctor with fever had a predicted value 3.3 (1.8-4.9) higher than others (p=0.0005).

Examining age patterns of seroprevalence, we found an increase in the proportion seropositive against DENV with age in both 2013 and 2015, and a rise in seroprevalence was observed in almost all age groups after the 2013/14 outbreak (Figure 3A). However, the high levels of seroprevalence in older age groups made it challenging to estimate age-specific probability of infection, because there was a relative lack of serologically naive individuals in these groups to act as a denominator (Table 4). We therefore again used rise in ELISA value as a correlate of infection, based on Figure 2B. As well as producing more precise estimates of infection risk in older groups (Table 4), this approach also suggested that individuals aged 10–19 years were most likely to be infected. This is in contrast to the surveillance data, which indicated the highest per capita level of reported disease was in the 20–29 age group (Figure 3B).

**Figure 3:**
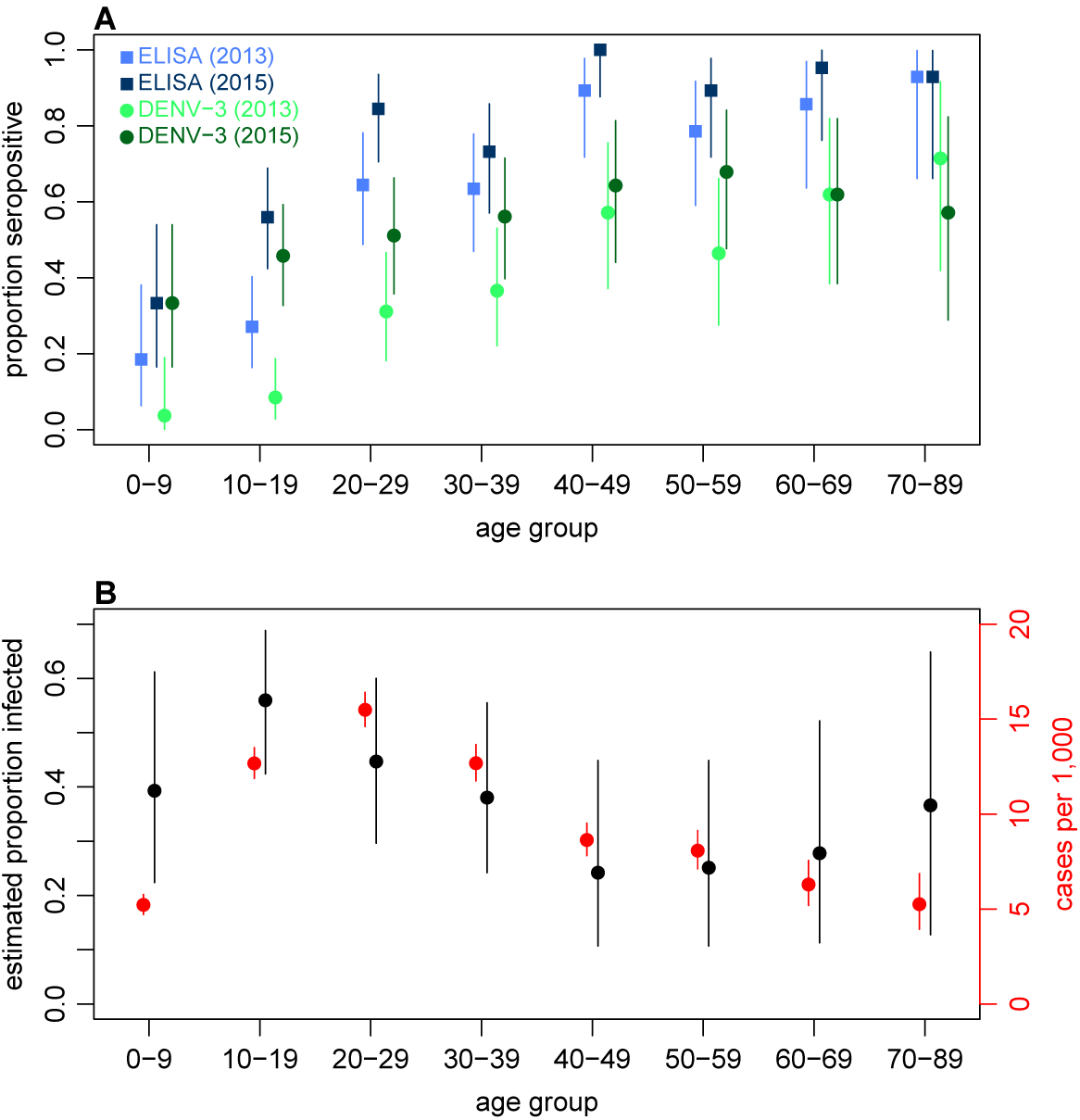
Age patterns of immunity and infection during 2013–15. (A) Proportion of each age group seropositive against DENV as measured by ELISA (blue squares) and DENV-3 by MIA (green circles). Lighter points show 2013 results, darker points show 2015; lines show 95% binomial confidence intervals. (B) Comparison of estimated age-specific infection and reported cases. Black points, estimated proportion infected based on ELISA rise indicated in Figure 2B; red points, cases reported per 1,000 people in each age group; lines show 95% binomial confidence intervals.

**Table 4:** Estimated age-specific attack rates based on adjusted distribution of ELISA values, and seroconversion using ELISA cutoff. Estimated number of infections were calculated from the total of the probabilities that each individual in that age group had been infected, to the nearest integer. Binomial 95% confidence intervals are shown in parentheses.

| Age | N | Infections | % | Seronegative | Seroconverted | % |
| --- | --- | --- | --- | --- | --- | --- |
| 0–9 | 27 | 10 | 37% (19.4-57.6%) | 22 | 6 | 27.3% (10.7-50.2%) |
| 10–19 | 59 | 30 | 50.8% (37.5-64.1%) | 43 | 17 | 39.5% (25-55.6%) |
| 20–29 | 45 | 18 | 40% (25.7-55.7%) | 16 | 10 | 62.5% (35.4-84.8%) |
| 30–39 | 41 | 13 | 31.7% (18.1-48.1%) | 15 | 6 | 40% (16.3-67.7%) |
| 40–49 | 28 | 6 | 21.4% (8.3-41%) | 3 | 3 | 100% (29.2-100%) |
| 50–59 | 28 | 6 | 21.4% (8.3-41%) | 6 | 3 | 50% (11.8-88.2%) |
| 60–69 | 21 | 4 | 19% (5.45-41.9%) | 3 | 3 | 100% (29.2-100%) |
| 70+ | 14 | 4 | 28.6% (8.39-58.1%) | 1 | 0 | 0% (0-97.5%) |
| Total | 263 | 89 | 33.8% (28.1-39.9%) | 109 | 48 | 44% (34.5-53.9%) |

Next, we explored spatial patterns of infection in different communities. Previous studies have suggested that dengue outbreaks can spread outwards from urban hubs to more rural areas [46, 47]. A similar spatial pattern was observed from the surveillance data during the early stages of the 2013/14 Fiji outbreak (Figure 4A). The first case was reported at Colonial War Memorial Hospital (CWM), Fiji’s largest hospital located in central urban Suva, in the week ending 4th November 2013. The outbreak took 9 weeks to reach the furthest reporting point from CWM in Central Division, a health centre 51km away by Euclidean distance (i.e. as the crow flies). We found limited association between Euclidean distance from CWM and proportion of study cluster seropositive to DENV-3 in 2015 (Figure 4B): the Pearson correlation between ELISA seropositivity in each cluster and distance from CWM was *ρ*= –0.12 (p=0.59); for DENV-3 the correlation coefficient was *ρ*= –0.46 (p=0.03). However, we found no significant association between the Euclidean distance from CWM and proportion of cluster infected (Figure 4C). Pearson correlation between estimated proportion infected based on change ELISA value in each cluster and distance from CWM was *ρ*= 0.22 (p=0.30); for DENV-3 the correlation was *ρ*= –0.36 (p=0.09). We did find evidence of dengue seroconversion in every cluster, however, suggesting that the outbreak eventually spread throughout Central Division.

**Figure 4:**
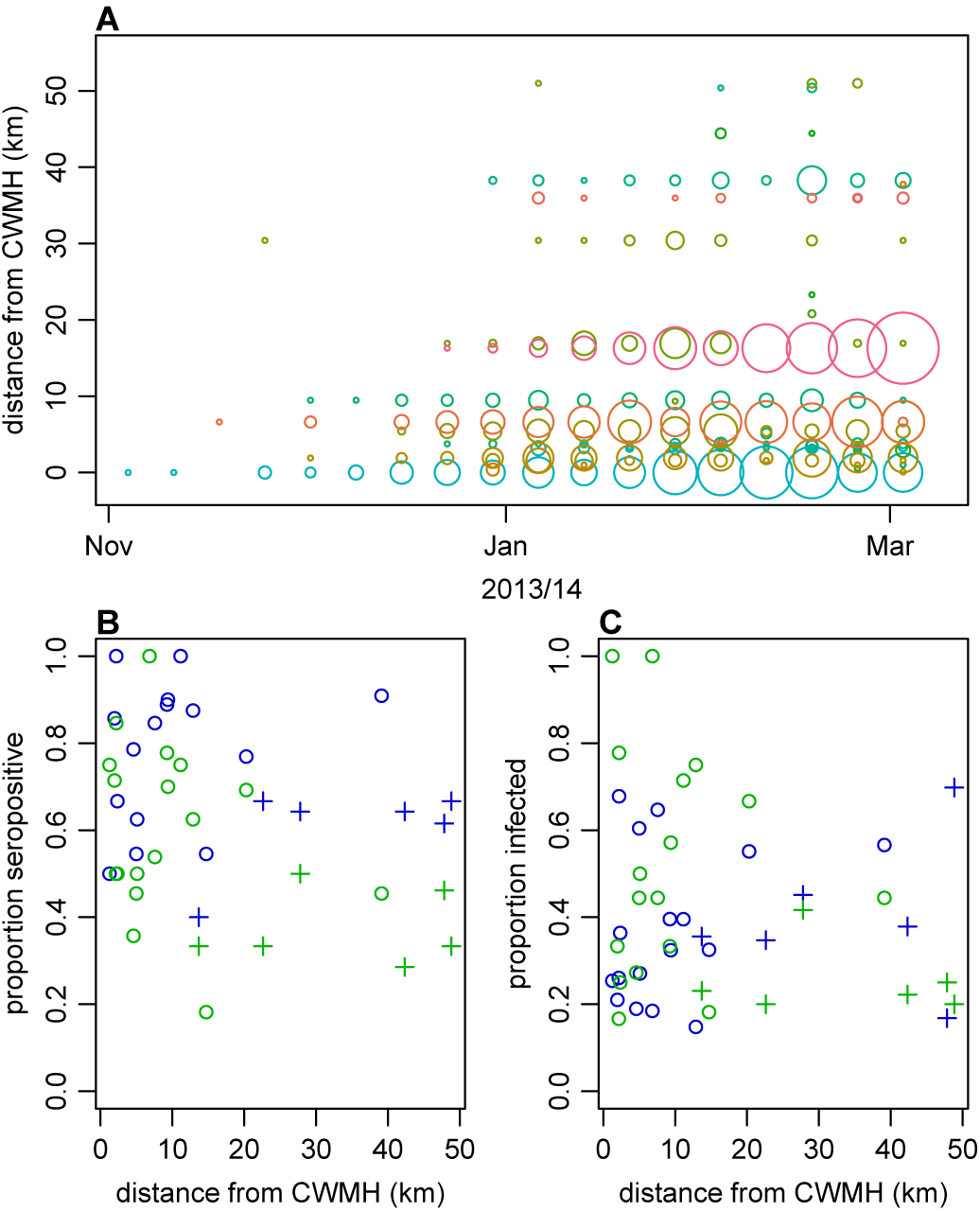
Spatial pattern of infection and immunity in Central Division. (A) Relationship between dengue cases reported by each health centre at the start of the outbreak and Euclidean distance from Colonial War Memorial Hospital (CWM) in Suva. Area of circle is proportional to number of cases reported in that week; each health centre is represented by a different colour. (B) Proportion seropositive in each serosurvey study cluster in 2015 vs Euclidean distance from CWM. Blue, ELISA data; green, MIA data; circles, urban or peri-urban clusters; crosses, rural clusters. (C) Proportion infected in each serosurvey study cluster vs Euclidean distance from CWM. Blue, estimate based on ELISA data, using adjustment in Figure 2B; green, seroconversion based on MIA for individuals who were initially seronegative; circles, urban or peri-urban clusters; crosses, rural clusters.

As we did not find strong individual or community-level heterogeneity in infection, we used both the surveillance data and paired serological survey to test explanations for the observed outbreak dynamics at the division level. Fitting an age-structured mathematical model of vector-borne infection dynamics to the surveillance data alone, we could reproduce the observed incidence pattern under the assumption of a simple immunising epidemic. Specifically, the reported cases were consistent with an epidemic that declined as a result of depletion of the susceptible population (Figure S6). However, this basic epidemic model underestimated initial immunity and overestimated final immunity, as measured by seropositivity to DENV-3 by MIA in individuals under and over 20 years old. When we jointly fitted to surveillance data and age-specific serology, the simple model could not reproduce both sets of data (Figure S7). A similar discrepancy between serological surveys and surveillance data has been noted in previous arbovirus modelling studies for French Polynesia and Micronesia [48, 18, 33].

The addition of seasonal variation in transmission improved model performance (Table S3), although the model suggested the outbreak lasted longer than the observed data suggested, with a predicted second peak in 2015 (Figure S8). To capture potential reduction in transmission following the introduction of a clean-up campaign in March 2014, we included a flexible additional sigmoidal transmission rate in the model, which was constrained so that the midpoint of the decline occurred after the start of the campaign on 8th March 2014. As well as performing better than the others tested as measured by deviance information criterion (DIC) (Table S3), this model was able to reproduce the observed surveillance and serological data (Figure 5A–B). The model indicated a reduction in transmission that coincided with the clean-up campaign (Figure 5C). As the effective reproduction number was near the critical value of one when the clean-up campaign was introduced (Figure 5D), it suggests that the main contribution of control measures may have been to bring DENV-3 infections to sufficiently low levels for transmission to cease, preventing persistence into the following season. We obtained the same conclusion when ELISA rather than MIA seroprevalence was used to quantify immunity during model fitting (Table S3 and Figure S9).

**Figure 5:**
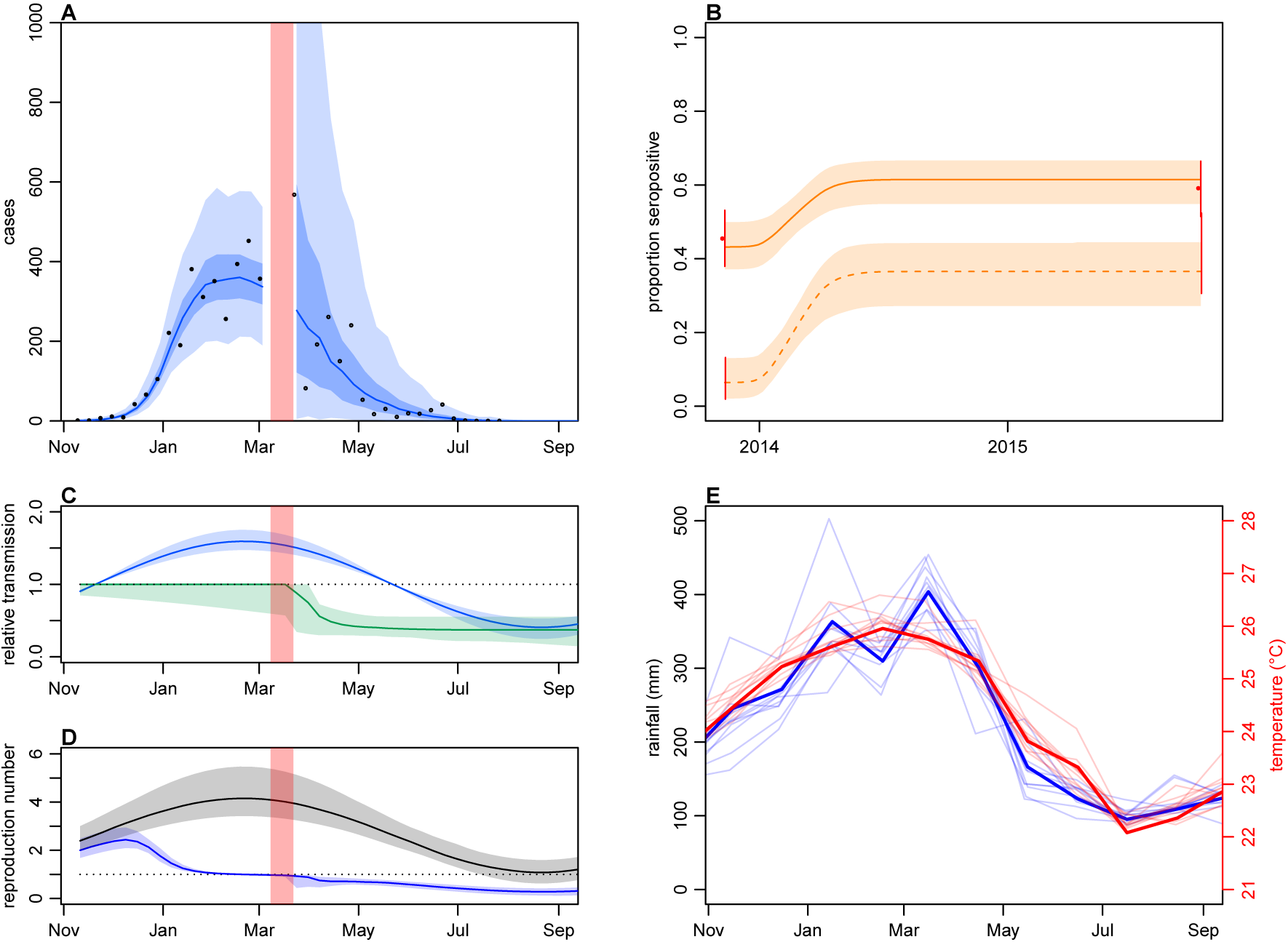
Impact of climate and control measures on DENV transmission during 2013/14, using model jointly fitted to surveillance and serological data. (A) Model fit to surveillance data. Solid black dots, confirmed dengue cases used for fitting model to early phase; black circles, DLI cases used for fitting late phase; blue line, median estimate from fitted model; dark blue region, 50% credible interval; light blue region, 95% CrI; red region shows timing of clean-up campaign. (B) Pre- and post-outbreak DENV immunity. Red dots show observed MIA seroprevalence against DENV-3 in autumn 2013 and autumn 2015; hollow dots, under 20 age group; solid dots, 20+ age group; lines show 95% binomial confidence interval. Dashed orange line shows model estimated rise in immunity during 2013/14 in under 20 group; solid line shows rise in 20+ group; shaded region shows 95% CrI. (C) Estimated variation in transmission over time. Red region, timing of clean-up campaign; blue line, relative transmission as a result of seasonal effects; green line, relative transmission as a result of control measures. Shaded regions show 95% CrIs. (D) Change in reproduction number over time. Black line, basic reproduction number, *R*_0_; blue line, effective reproduction number, *R*. Shaded regions show 95% CrIs. Dashed line shows the *R* = 1 herd immunity threshold. (E) Average monthly rainfall and temperature in Fiji between 2003–14; thick lines show data for 2013/14 season.

Fitting to the DENV-3 MIA seroprevalence data, we estimated that the mean basic reproduction number, *R*_0_, over the course of the year was 2.64 (95% CrI: 2.13–3.46), with a peak value of 4.15 (3.42–5.47) in February (Table 5). Accounting for under-reporting and stochastic variability in weekly case reporting, we estimated that 6.8% (3.8–12%) of infections were reported as laboratory-confirmed cases during the early phase of the outbreak and 5.8% (0.22–44%) were reported as suspected cases in the later stages.

**Table 5:** Parameter estimates for the 2013/14 dengue epidemic when the model was fitted to MIA or ELISA data. Median estimates are shown, with 95% credible intervals shown in parentheses. Mean *R*_0_ is the average basic reproduction number over a year; seasonal reduction measures the ratio between *R*_0_ at the peak and lowest temperature. Group-specific effective reproduction numbers at the onset of the outbreak are also shown: from mosquitoes to humans (*R*_*hv*_) and from humans to mosquitoes (*R*_*vh*_). Proportion reported was calculated by sampling from the negative binomial distribution that defines the model observation process (i.e. the credible interval reflects both underreporting and dispersion in weekly case reporting). 
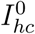
 and 
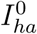
 denote the number of initially infectious individuals in the younger and older age group respectively.

| Parameter | MIA | ELISA |
| --- | --- | --- |
| Mean $R_0$ | 2.64 (2.13-3.46) | 3.27 (2.68-4.02) |
| Peak $R_0$ | 4.15 (3.42-5.47) | 5.18 (4.4-6.48) |
| Seasonal reduction | 0.744 (0.642-0.858) | 0.749 (0.643-0.886) |
| Control reduction | 0.628 (0.448-0.9737) | 0.585 (0.387-0.685) |
| $R_{hv}$ | 0.0139 (0.0084-0.0209) | 0.0122 (0.00801-0.0214) |
| $R_{vh}$ | 278 (162-574) | 405 (213-668) |
| Proportion reported, lab (%) | 6.8 (3.8-12) | 9.5 (5.1-19) |
| Proportion reported, suspected (%) | 5.8 (0.22-44) | 8.3 (0.55-52) |
| $I_{hc}(0)$ | 2.6 (1.1-21) | 2.4 (1.1-12) |
| $I_{ha}(0)$ | 6.6 (1.2-35) | 13 (3.3-39) |

## Discussion

We analysed surveillance reports and serological survey data to examine the dynamics of a major 2013/14 dengue outbreak in Fiji. Owing to the sporadic and unpredictable nature of dengue outbreaks in the Pacific [7], it is rare to have access to paired population-representative sera collected before and after such an epidemic. Comparing surveillance and serological survey data made it possible to understand the relationship between observed reported cases and the true attack rate and quantify the relative role of climate, herd immunity and control measures in shaping transmission.

Analysis of detailed serological data provided insights into age specific-patterns of infection that would not be identified from seropositivity thresholds alone. We estimated the highest infection rate was in the 10-19 year old age group, whereas proportionally the most reported cases were in the 20-29 year old group. The apparent disparity between reported cases and infections estimated from the serological survey may be the result of secondary DENV infections causing more severe clinical disease and therefore increasing the likelihood of seeking medical care [49]. The ELISA results suggested that fewer than 50% of individuals under age 20 had experienced DENV infection in 2013 (Figure 3A), which means an infection during the 2013/14 outbreak in this group was more likely to be primary than secondary. In contrast, the majority of 20–29 year olds already had evidence of infection in 2013, and hence 2013/14 outbreak would have generated relatively more secondary or tertiary infections in this group. In addition, if age-specific infection rates are indeed higher in younger groups, it means that estimating population attack rates based on the proportion of seronegative individuals infected may over-estimate the true extent of infection. Focusing on the seronegative subset of the population leads to children being over-sampled, which in our data inflates attack rate estimates by around 10% compared to estimates based on change in ELISA value (Table 4).

We also found little evidence of spatial heterogeneity in seroconversion. Although the locations of health centres reporting cases in the early stages of the outbreak suggested infection spread outwards from central Suva, we found evidence of DENV infection in all study clusters. This suggests that spatial structure may be more important in driving transmission dynamics early in the outbreak, but might not influence the final attack rate.

Analysis of risk factors found that presence of self-reported symptoms between 2013–15 was associated with DENV infection. There was also a strong association between rise in ELISA value and self-reported symptoms in individuals who were likely infected, which suggests that raw values from serological tests could potentially be used to estimate the proportion of a population who were asymptomatic during a dengue outbreak, even in older age groups that were already seropositive. However, it is worth noting that the questionnaire that accompanied the serosurvey was brief and only asked about fever and visits to a doctor with fever; there may be specific factors that can better predict prior infection in such settings. We also conducted the follow up survey around 18 months after the outbreak, which means recall bias is a potential limitation of the risk factor analysis.

To investigate potential explanations for the outbreak decline in early 2014, we fitted transmission dynamic models to both surveillance and serological survey data. Our analysis shows the benefits of combining multiple data sources: with surveillance data alone, it would not have been possible to distinguish between self-limiting outbreak driven by a decline in the susceptible population, and one that had ceased for another reason. With the addition of serological data in the model fitting, however, we were able to quantify the relative contribution of herd immunity, climate and control measures to the outbreak dynamics. In particular, seasonal variation and herd immunity alone could fully explain the fall in transmission: the seasonal model predicted a second outbreak wave in early 2015, which did not occur in reality. However, an additional decline in transmission in March 2014, which coincided with a nationwide vector clean-up campaign, could capture the observed patterns in serological and surveillance data, and in the model prevented a second wave of DENV-3 infections.

There are some limitations to our modelling analysis. First, we assumed that seropositivity in IgG antibody tests was equivalent to protective immunity. High levels of neutralising antibodies have been shown to correlate with protection from symptomatic infection [50], but it remains unclear precisely how much an individual with a given ELISA or MIA value contributes to transmission. Second, we focused on seroprevalence against DENV-3 in the main modelling analysis. As prior infection with one dengue serotype can lead to a cross-reactive immune response against other serotypes [6], we fitted the model to ELISA data (which is not serotype specific) as a validation; this produced the same overall conclusions. Third, we used a flexible time-dependent transmission rate to capture a potential reduction in transmission as a result of control measures in March 2014. The clean-up campaign included multiple concurrent interventions, which occurred alongside ongoing media coverage of the outbreak; it was therefore not possible to untangle how specific actions – such as vector habitat removal or changes in community behaviour that reduced chances of being bitten – contributed to the outbreak decline. Moreover, factors unrelated to control, such as spatial structure or local weather effects, may also have contributed to the observed decline in transmission; there was heavy rain and flooding in Viti Levu at the end of February 2014 [51]. Finally, our analysis focused on Central Division, Fiji. However, much of the data used in our model – such as surveillance data, post-outbreak serology, and climate – would be available for other settings. For factors that are harder to measure without paired serology, like age-specific infection rates and potential effectiveness of control measures, a joint inference approach could be employed that combines prior distributions based on the data presented here with available outbreak data from the other location of interest [18].

Despite these caveats, our results show that transmission dynamic models developed using a combination of serological surveys and surveillance data, can be valuable tool for examining dengue fever outbreaks. As well as providing insights into the transmission and control of dengue, the analysis has implications for forecasting of future epidemics. During February and March 2014, members of the research team based at London School of Hygiene & Tropical Medicine provided real-time analysis and outbreak projections for the Fiji National Centre for Communicable Disease Control, to support public health planning [52]. However, a lack of serological data at the time meant it was necessary to make strong assumptions about pre-existing population immunity. With up-to-date population representative serology now available, forecasting models during future outbreaks will be able to include a more realistic herd immunity profile from the outset. Such seroepidemiological approaches could also be employed in other settings, to provide improved forecasts of dengue transmission dynamics and potential disease burden prior to and during outbreaks, as well as quantitative retrospective evaluation of the effectiveness of control measures.

## Acknowledgements

AWe warmly thank all the participants and community leaders who generously contributed to the study. We are also grateful to Kylie Jenkins of Australian Aid’s Fiji Health Sector Support Programme, Teheipuaura Mariteragi-Helle at the Institut Louis Malardé, and Dr Ketan Christie at the University of the South Pacific. We thank the staff of the Ministry of Health clinical services and Mataika House Fiji Centre for Communicable Disease Control for providing the surveillance data underpinning this study. We would also like to acknowledge the work of the field teams: Dr. Kitione Rawalai, Jeremaia Coriakula, Ilai Koro, Sala Ratulevu, Ala Salesi, Meredani Taufa, and Leone Vunileba (2013); Meredani Taufa, Adi Kuini Kadi, Jokaveti Vubaya, Colin Michel, Mereani Koroi, Atu Vesikula, and Josateki Raibevu (2015).

## Author contributions

AJK, MK, CHW, CLL, WJE, JA, EJN, VMCL, SH and MH contributed to study design. AJK and CHW coordinated the 2015 sample collection. JvH and JCM developed the MIA protocol. MK, MA, JA and VMCL undertook sample testing. AJK performed statistical and mathematical modelling. AJK, MK, CHW, MA, SF, ADH, OJB, WJE, EJN, VMCL, SH, MH reviewed and interpreted results. AJK and CHW wrote the first draft of the manuscript. AJK, MK, CHW, MA, SF, OJB, CLL, JA, EJN, VMCL, SH, MH commented on and edited draft versions of the manuscript and all authors approved the manuscript.

## S1 Supporting Information

**Table S1:** Relationship between seroconversion and health-seeking behaviour for participants who were initially seronegative by ELISA in 2013. Table shows breakdown for participants who were also negative in 2015 (i.e. did not seroconvert) and positive in 2015 (i.e. seroconverted).

| Health-seeking behaviour | Negative | % (95% CI) | Positive | % (95% CI) |
| --- | --- | --- | --- | --- |
| No fever in preceding two years | 77 | 79.4 (70-86.9) | 26 | 68.4 (51.3-82.5) |
| Fever in preceding two years without visiting doctor | 4 | 4.12 (1.13-10.2) | 2 | 5.26 (0.644-17.7) |
| Visited doctor with fever in preceding two years | 16 | 16.5 (9.73-25.4) | 10 | 26.3 (13.4-43.1) |
| Total | 97 |  | 38 |  |

**Table S2:** Parameters used in the model. Prior distributions are given for all parameters, along with source if the prior incorporates a specific mean value. All rates are given in units of days^−1^.

| Parameter | Definition | Prior | Source |
| --- | --- | --- | --- |
| $1/\alpha_v$ | extrinsic latent period | Gamma($\mu=15$, $\sigma=0.1$) | [59] |
| $1/\alpha_h$ | intrinsic latent period | Gamma($\mu=5.9$, $\sigma=0.1$) | [59] |
| $1/\gamma$ | human infectious period | Gamma($\mu=5$, $\sigma=0.1$) | [60] |
| $1/\delta$ | mosquito lifespan | Gamma($\mu=8$, $\sigma=0.1$) | [61] |
| $\hat{\beta}$ | baseline vector-to-human transmission rate | $\mathcal{U}(0, \infty)$ | |
| $a_v$ | relative human-to-vector transmission | $\mathcal{U}(0, \infty)$ | |
| $a_0$ | amplitude of seasonal transmission | Gamma($\mu=0.592$, $\sigma=0.01$) | [38, 39] |
| $a_1$ | gradient of sigmoidal change in transmission | $\mathcal{U}(0, \infty)$ | |
| $a_2$ | magnitude of sigmoidal change in transmission | $\mathcal{U}(0, 1)$ | |
| $a_\tau$ | timing of sigmoidal change in transmission | $\mathcal{U}(8\text{th March } 2014, 5\text{th April } 2014)$ | [11] |
| $r_L$ | proportion of cases reported as lab confirmed | $\mathcal{U}(0, 1)$ | |
| $\phi_L$ | reporting dispersion, lab | $\mathcal{U}(0, \infty)$ | |
| $r_S$ | proportion of cases reported as suspected | $\mathcal{U}(0, 1)$ | |
| $\phi_S$ | reporting dispersion, suspected | $\mathcal{U}(0, \infty)$ | |
| $I_{hc}^0$ | initial number infectious aged <20 | $\mathcal{U}(0, N_c)$ | |
| $R_{hc}^0$ | initial number immune aged <20 | $\mathcal{U}(0, N_c)$ | |
| $I_{ha}^0$ | initial number infectious aged 20+ | $\mathcal{U}(0, N_a)$ | |
| $R_{ha}^0$ | initial number immune aged 20+ | $\mathcal{U}(0, N_a)$ | |
| $I_v^0$ | initial proportion of infectious mosquitoes | $\mathcal{U}(0, 1)$ | |

**Table S3:** Comparison of model performance. Gap denotes the number of weeks between the end of the first phase of data fitting (i.e. fitting to confirmed cases) and the start of the second phase of fitting to DLI cases.

| Model | Serological data | Gap (weeks) | DIC | ΔDIC |
| --- | --- | --- | --- | --- |
| SEIR | MIA | 2 | 713.7 | 358.1 |
| SEIR + climate | MIA | 2 | 376.2 | 20.59 |
| SEIR + climate + control | MIA | 2 | 355.6 | 0 |
| SEIR | MIA | 3 | 7412 | 7078 |
| SEIR + climate | MIA | 3 | 357.1 | 23.04 |
| SEIR + climate + control | MIA | 3 | 334 | 0 |
| SEIR | MIA | 4 | 368.9 | 43.52 |
| SEIR + climate | MIA | 4 | 383.1 | 57.69 |
| SEIR + climate + control | MIA | 4 | 325.4 | 0 |
| SEIR | ELISA | 2 | 2852 | 2488 |
| SEIR + climate | ELISA | 2 | 369.2 | 5.301 |
| SEIR + climate + control | ELISA | 2 | 363.9 | 0 |

**Figure S1:**
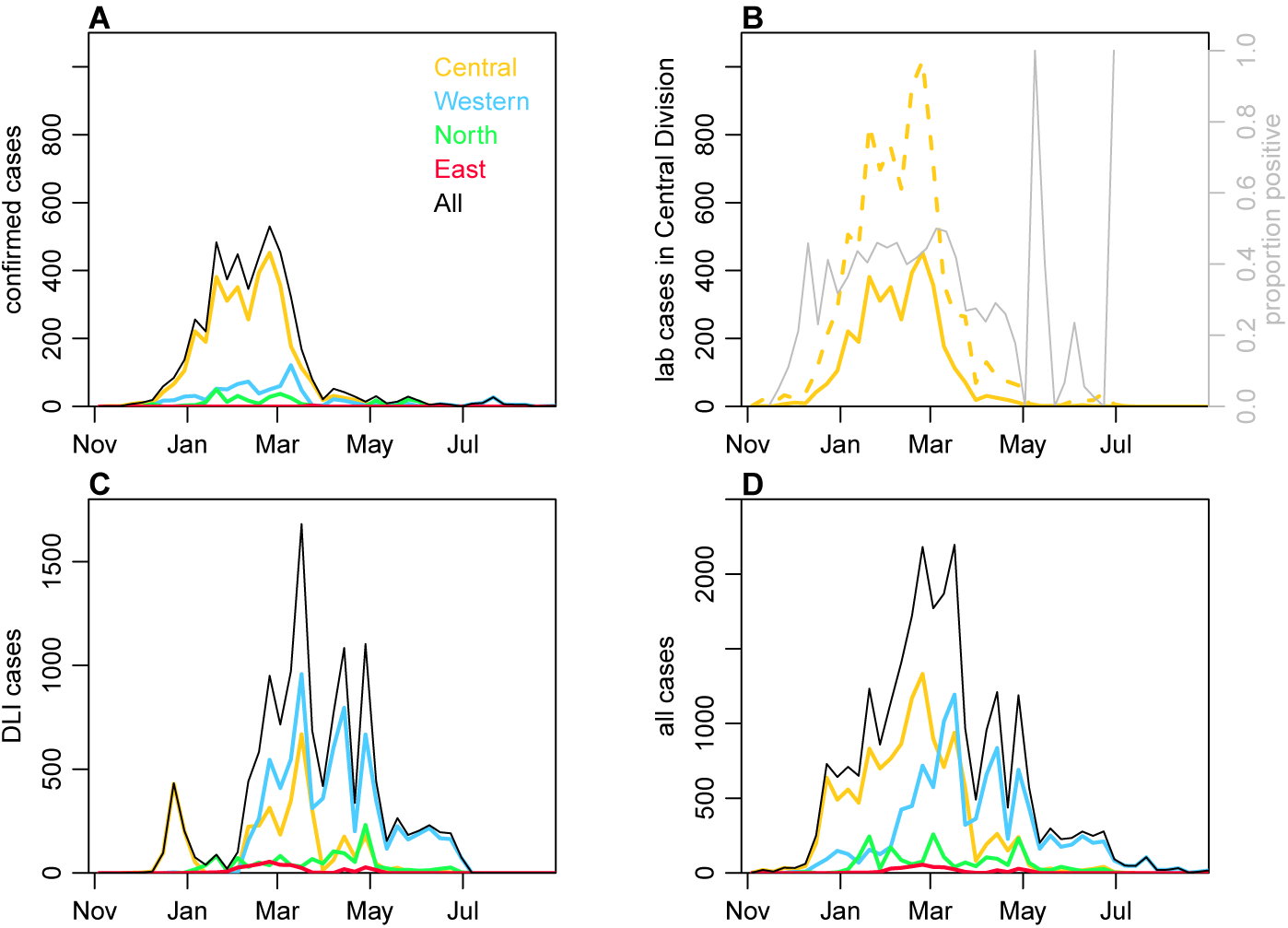
Geographical distribution of cases. (A) Lab confirmed dengue cases reported in Northern (green), Western (blue) and Central (yellow) divisions between 27th October 2013 and 31st August 2014. (B) Total tested and confirmed cases in Central division (dashed and solid lines respectively), as well as proportion of cases that tested positive (grey line). (C) Dengue-like illness (DLI) over time. (D) Total suspected cases (i.e. tested and DLI).

**Figure S2:**
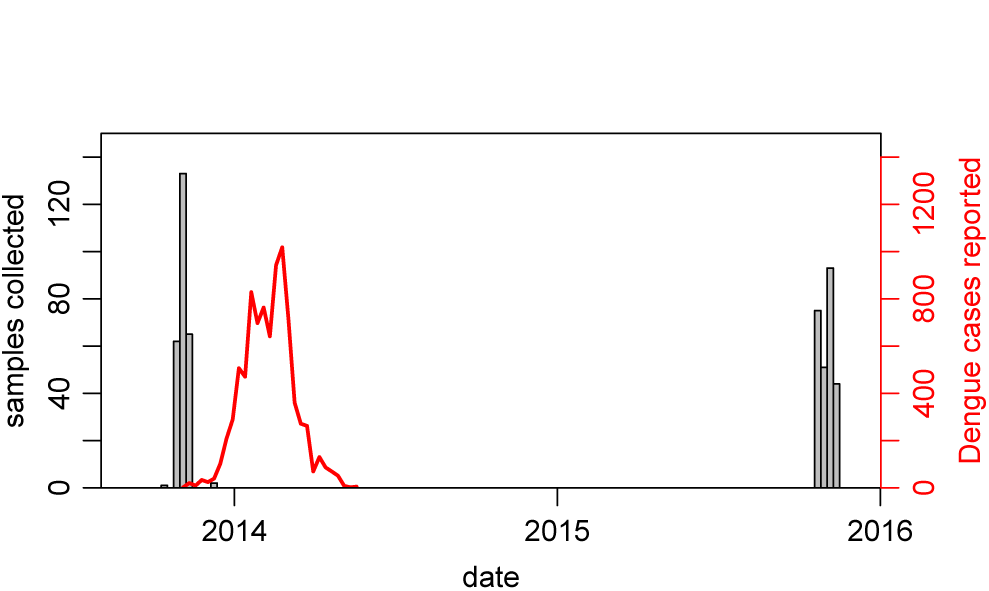
Dates of serological sample collection. Grey bars show weekly number of samples collected in Central Division across the two studies (paired samples shown only). Red line shows lab tested cases in Central Division.

**Figure S3:**
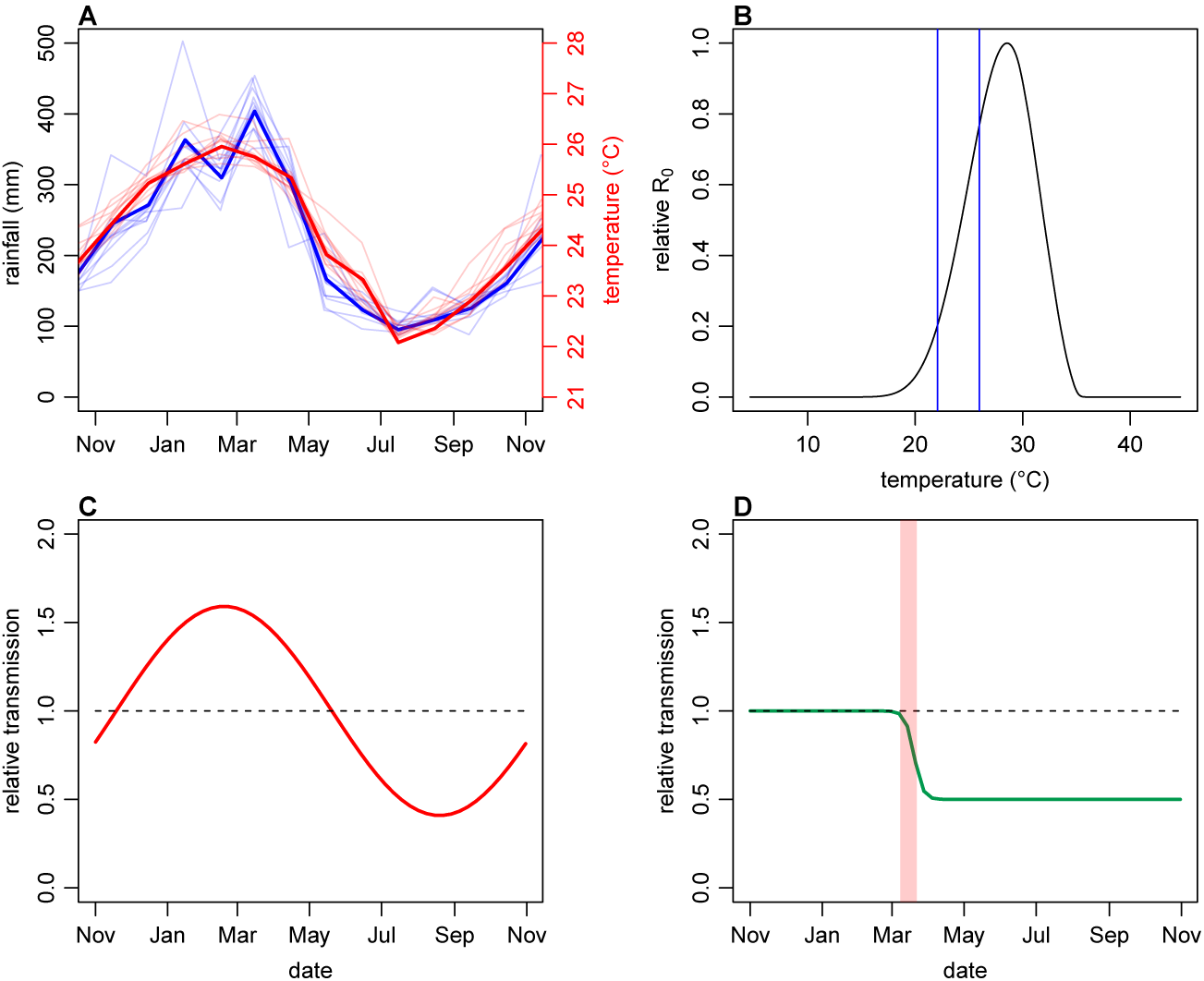
Illustration of model variation in transmission as a result of climate and control. (A) Observed rainfall and temperature in Fiji between 2003–14; thick lines show 2013/14 season. (B) Relationship between temperature and relative basic reproduction number *R*_0_ (adapted from [39]). Blue lines show maximum and minimum temperature observed in Fiji during the 2013/14 season. (C) Expected seasonal variation transmission based on the expected decline between the blue lines in (B). Seasonality following a sine wave that is shifted so that it reaches its maximum at the same time as the observed Fiji climate data. (D) Illustrative example of a sigmoidal drop in transmission after clean-up campaign introduced on 8th March 2014. Here we assume a decline of 50%; in the model analysis this parameter is fitted, along with the gradient of the decline.

**Figure S4:**
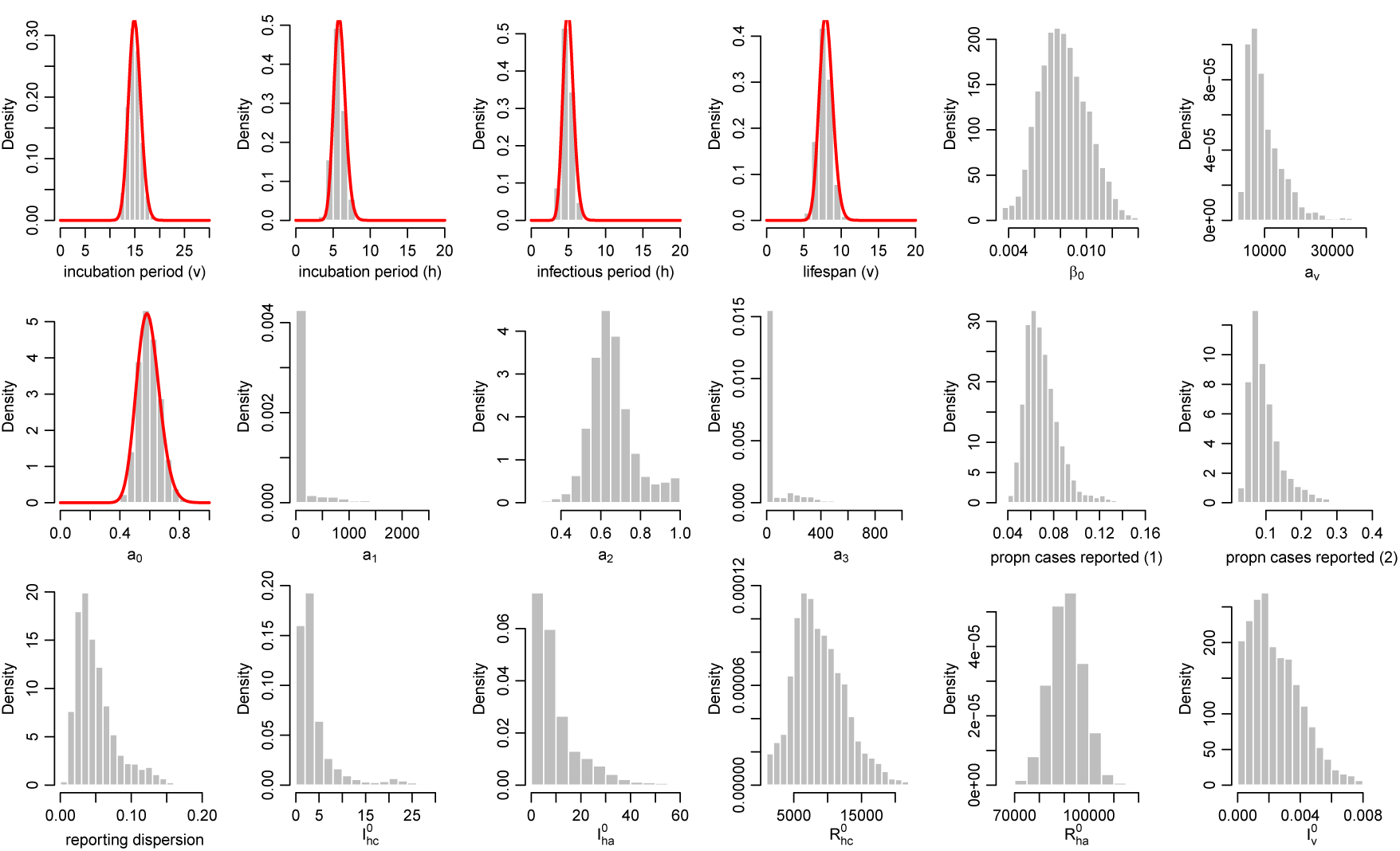
Posterior parameter estimates for the model with climate and control measures. Histograms show the estimated posterior distributed from the MCMC chain, discarding burn in iterations, for each parameter in Table S2. Red lines show prior distributions if informative priors were used for that parameter.

**Figure S5:**
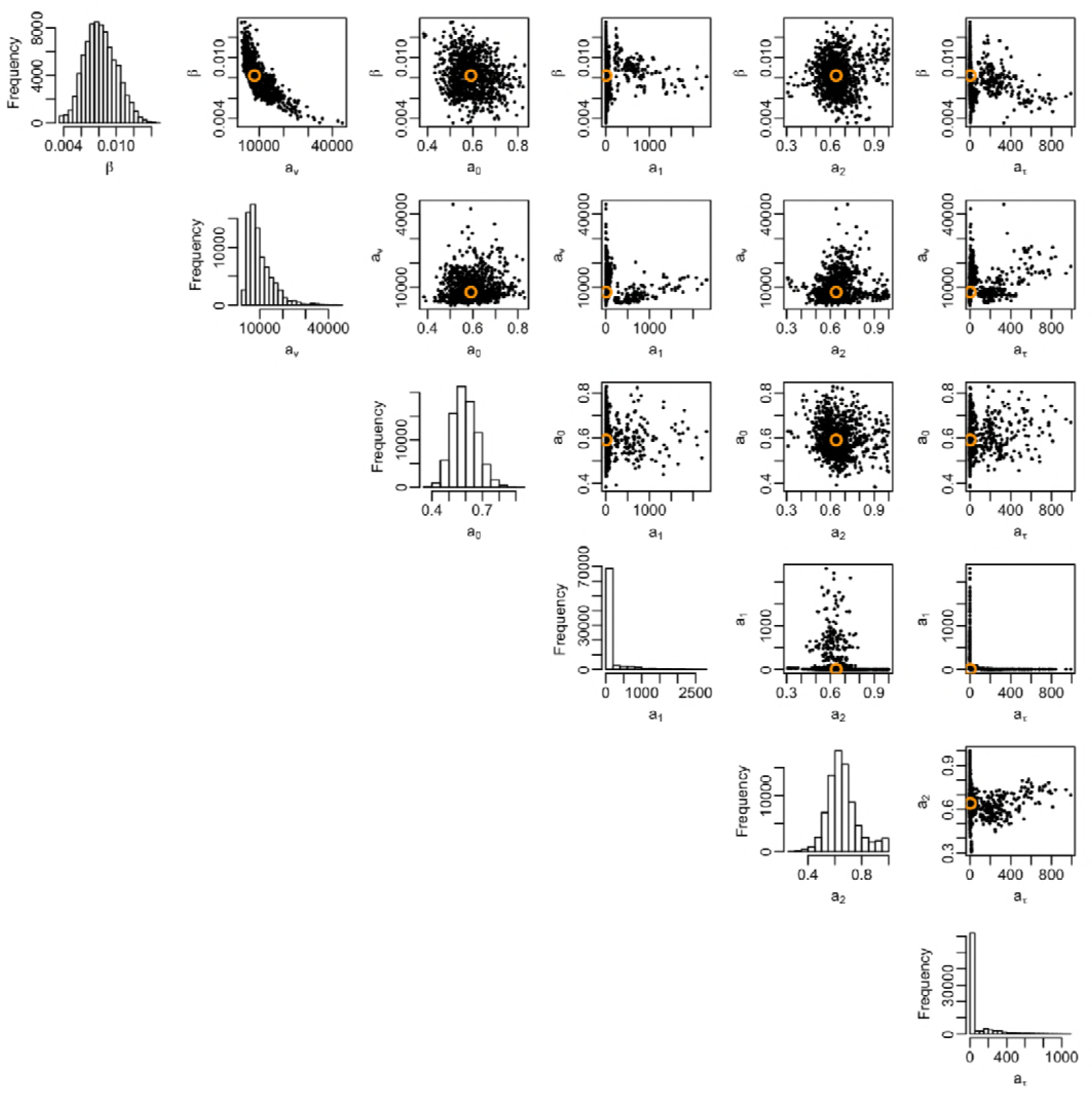
Correlation between posterior distributions of transmission rate parameters. Black dots show samples from the joint posterior distribution, with median given by orange circle. Histograms show the marginal posterior for each parameter.

**Figure S6:**
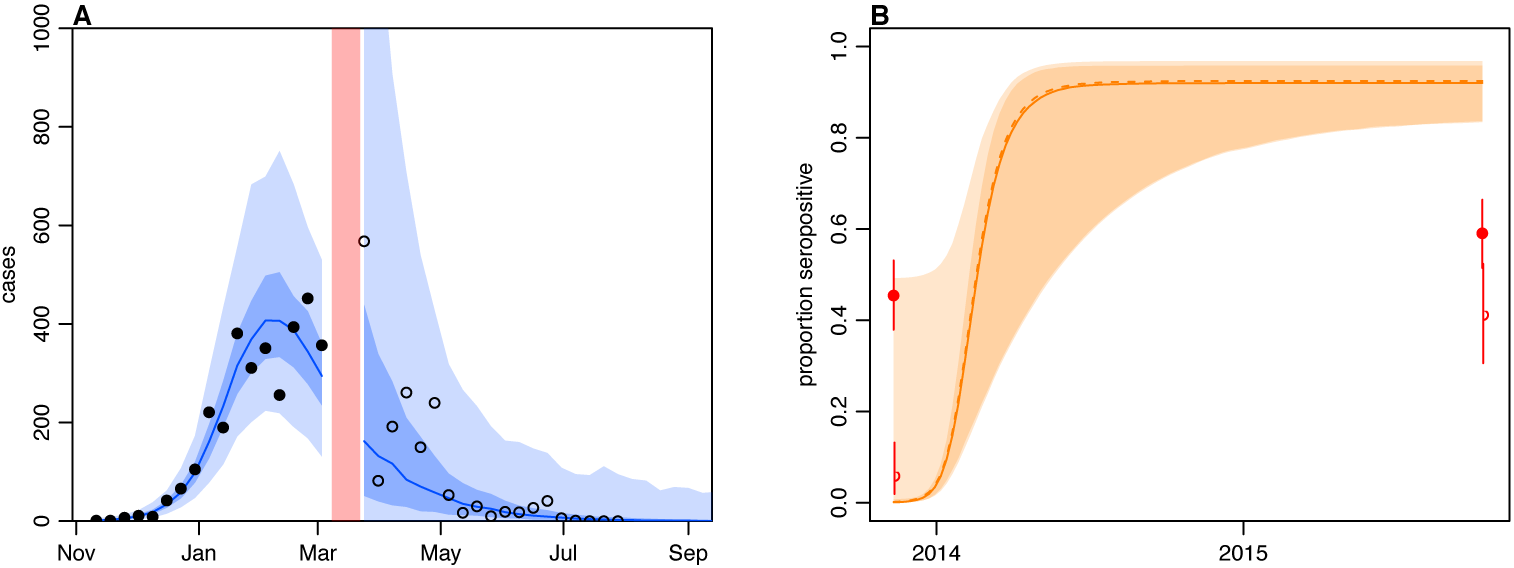
Dynamics of model fitted to case data only, without time-varying transmission. (A) Model fit to surveillance data. Black dots, suspected dengue cases; blue line, median estimate from fitted model; dark blue region, 50% credible interval; light blue region, 95% CrI; red region shows timing of clean-up campaign. (B) Pre- and post-outbreak DENV immunity. Red dots show observed MIA seroprevalence against DENV-3 in autumn 2013 and autumn 2015; hollow dots, under 20 age group; solid dots, 20+ age group; lines show 95% binomial confidence interval. Dashed orange line shows model estimated rise in immunity during 2013/14 in under 20 group; sold line shows rise in 20+ group; shaded region shows 95% CrI.

**Figure S7:**
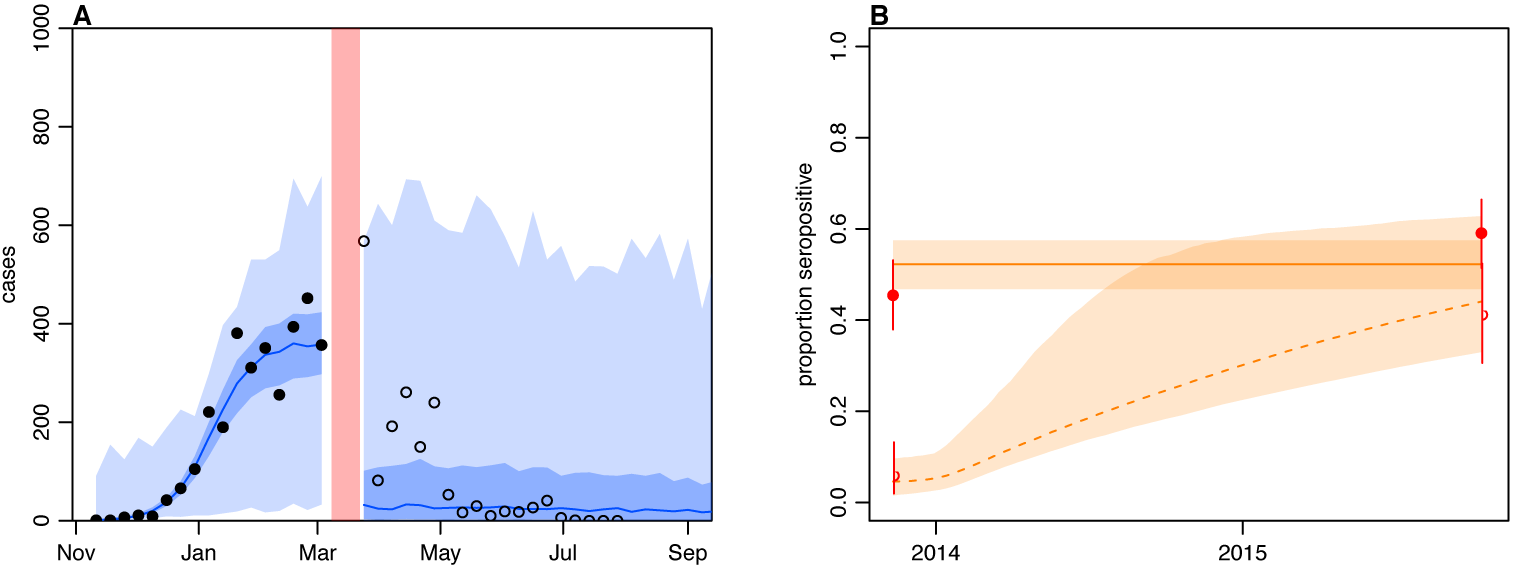
Dynamics of model jointly fitted to surveillance and serological data, without time-varying transmission. (A) Model fit to surveillance data. Black dots, suspected dengue cases; blue line, median estimate from fitted model; dark blue region, 50% credible interval; light blue region, 95% CrI; red region shows timing of clean-up campaign (B) Pre- and post-outbreak DENV immunity. Red dots show observed MIA seroprevalence against DENV-3 in autumn 2013 and autumn 2015; hollow dots, under 20 age group; solid dots, 20+ age group; lines show 95% binomial confidence interval. Dashed orange line shows model estimated rise in immunity during 2013/14 in under 20 group; sold line shows rise in 20+ group; shaded region shows 95% CrI.

**Figure S8:**
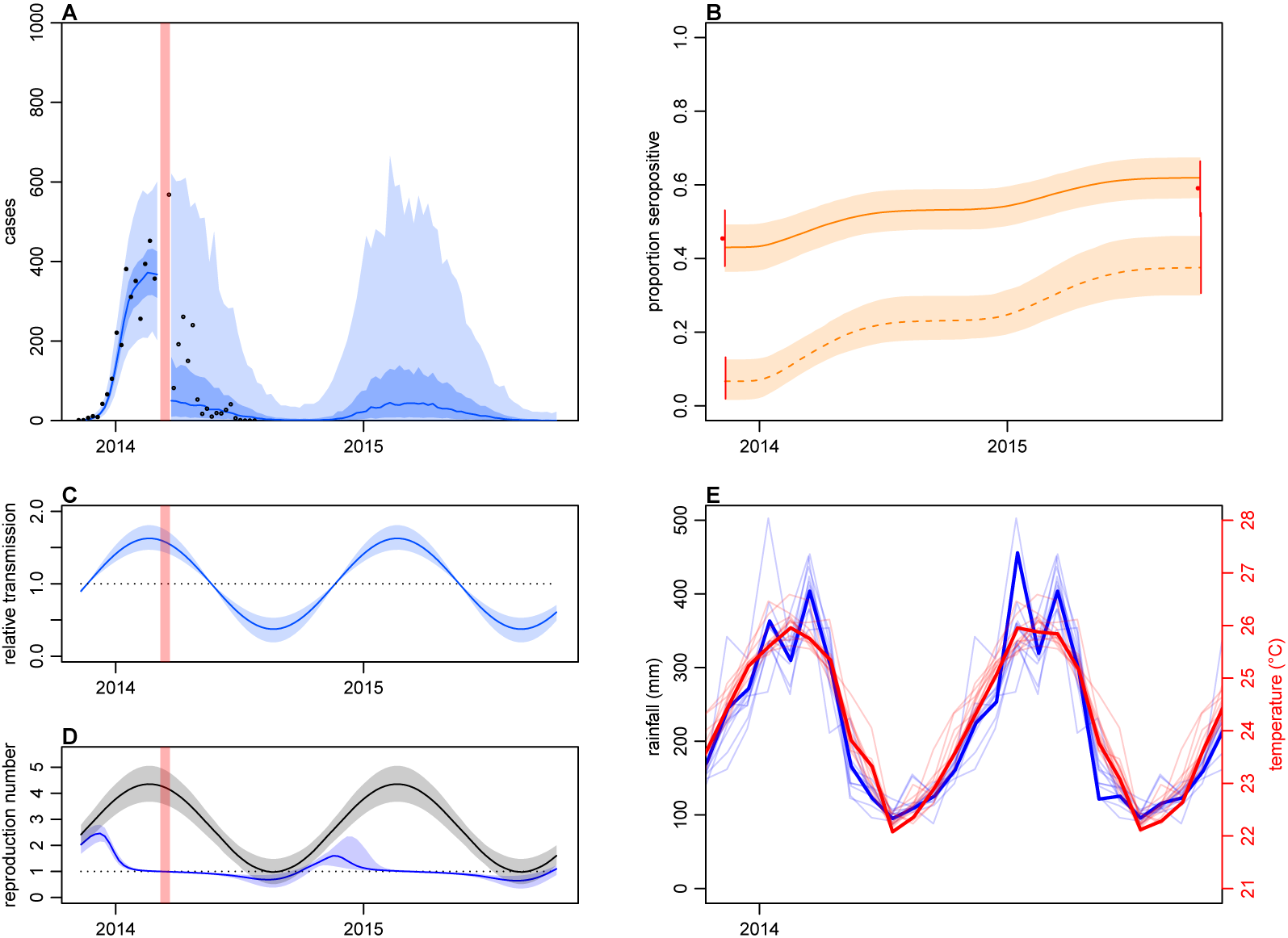
Dynamics of model jointly fitted to surveillance and serological data, with only climate-based variation in transmission. (A) Model fit to surveillance data. Solid black dots, confirmed dengue cases used for fitting model to early phase; black circles, DLI cases used for fitting late phase; blue line, median estimate from fitted model; dark blue region, 50% credible interval; light blue region, 95% CrI; red region shows timing of clean-up campaign. (B) Pre- and post-outbreak DENV immunity. Red dots show observed MIA seroprevalence against DENV-3 in autumn 2013 and autumn 2015; hollow dots, under 20 age group; solid dots, 20+ age group; lines show 95% binomial confidence interval. Dashed orange line shows model estimated rise in immunity during 2013/14 in under 20 group; solid line shows rise in 20+ group; shaded region shows 95% CrI. (C) Estimated variation in transmission over time. Red region, timing of clean-up campaign; blue line, relative transmission as a result of seasonal effects; green line, relative transmission as a result of control measures. Shaded regions show 95% CrIs. (D) Change in reproduction number over time. Black line, basic reproduction number, *R*_0_; blue line, effective reproduction number, R. Shaded regions show 95% CrIs. Dashed line shows the *R* = 1 herd immunity threshold. (E) Average monthly rainfall and temperature in Fiji between 2003–14; thick lines show data for 2013/14 season.

**Figure S9:**
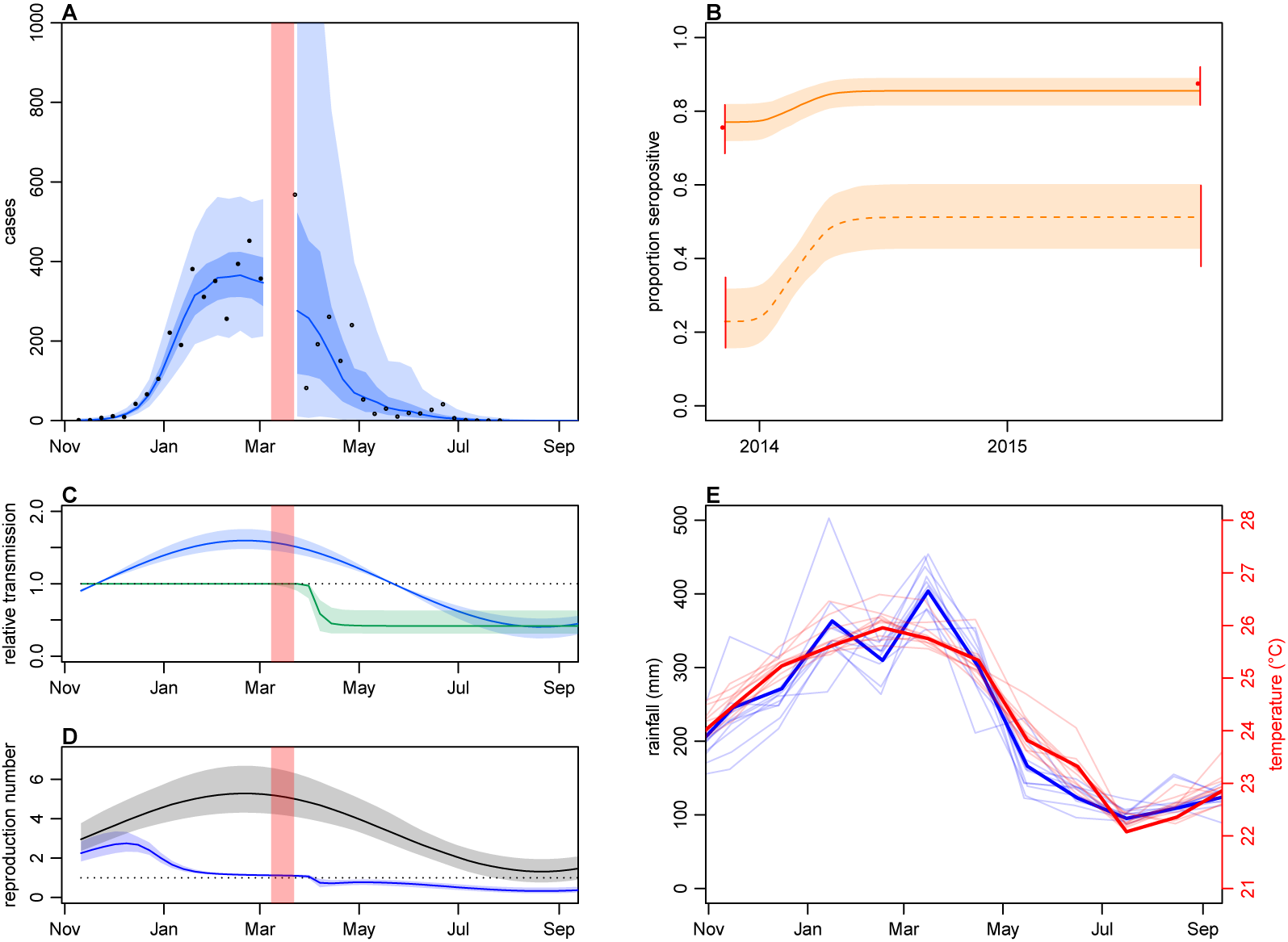
Impact of climate and control measures on dengue transmission during 2013/14, using model jointly fitted to surveillance and ELISA serological data. (A) Model fit to surveillance data. Solid black dots, confirmed dengue cases used for fitting model to early phase; black circles, DLI cases used for fitting late phase; blue line, median estimate from fitted model; dark blue region, 50% credible interval; light blue region, 95% CrI; red region shows timing of clean-up campaign. (B) Pre- and post-outbreak DENV immunity. Red dots show observed ELISA seroprevalence against DENV in autumn 2013 and autumn 2015; hollow dots, under 20 age group; solid dots, 20+ age group; lines show 95% binomial confidence interval. Dashed orange line shows model estimated rise in immunity during 2013/14 in under 20 group; solid line shows rise in 20+ group; shaded region shows 95% CrI. (C) Estimated variation in transmission over time. Red region, timing of clean-up campaign; blue line, relative transmission as a result of seasonal effects; green line, relative transmission as a result of control measures. Shaded regions show 95% CrIs. (D) Change in reproduction number over time. Black line, basic reproduction number, *R*_0_; blue line, effective reproduction number, R. Shaded regions show 95% CrIs. Dashed line shows the *R* = 1 herd immunity threshold. (E) Average monthly rainfall and temperature in Fiji between 2003–14; thick lines show data for 2013/14 season.

